# Early Steps in Pathogen-induced TTR-amyloid-formation

**DOI:** 10.64898/2026.09.21.753243

**Authors:** Malinda B. Premathilaka, Ulrich H. E. Hansmann

## Abstract

Motivated by correlations between SARS-COV-2 infections and transthyretin (TTR) amyloidosis we use molecular dynamics simulations to study whether three protein fragments from the SARS-CoV-2 virus can initiate dissociation of the transthyretin (TTR) homotetramer as the first step on the pathway to TTR amyloidosis. We find that the peptides reduce the frequency of transient hydrogen bonds at the monomer-monomer interfaces, decreasing in this way the stability of the TTR tetramer.

## Introduction

A hallmark of many neurodegenerative, metabolic and other diseases is the presence of amyloids deposits, but the mechanism of their formation and propagation into fibrils is only partially understood.^1^ Various *in vitro* studies suggest that microbial proteins and protein fragments can promote formation of amyloids of Aβ, α-synuclein (αS), amylin (type II diabetes) or Serum Amyloid A.^2–8^ Most of the previous research focused on their ability to modulate the aggregation of these proteins, encouraging formation of amyloids or stabilizing the resulting oligomers and fibrils.^3–6^ However, in many cases the crucial step may be instead the pathogen-induced unfolding of the native (i.e., functional) conformation of a protein that subsequently aggregates into amyloids. Pathogen induced misfolding of native conformation has been previously reported for prion protein and serum amyloid A.^9,10^

Another example for such a scenario could be transthyretin (TTR). This homotetrameric protein is expressed in the liver and choroid plexus and transports thyroxine and the retinol-binding protein.^11–14^ When the tetramer dissociates, the released monomers can partially unfold and aggregate into insoluble fibrils that may damage a wide spectrum of organs.^11,12,15–18^ For instance, their accumulation in the heart impedes normal cardiac function, and such TTR cardiac amyloidosis is seen in about 20% of elderly patients with heart failure.^19,20^ Drugs such as Vyndaqel (tafamidis meglumine) aim to stabilize the tetramer to reduce the number of chains that could potentially misfold and aggregate, in this way reducing the progression of fibril formation.^21–23^ As correlations between TTR amyloidosis and SARS-CoV-2 infections have been observed,^24^ we conjecture that SARS-CoV-2 derived proteins or protein fragments can induce TTR amyloidosis by an opposite effect, namely by de-stabilizing the tetramer.

In the present study we explore the plausibility of this hypothesis through large-scale all-atom molecular dynamics simulations. We consider three SARS-CoV-2 protein fragments: the protein fragments FI10 (^194^FKNIDGYFKI^203^) and HV11 (^1058^HGVVFLHVTYV^1068^), cleaved by the enzyme neutrophil elastase from the spike protein,^25^ and the segment SK9 (^55^SFYVYSRVK^63^) of the envelope protein with which we have worked extensively in the past.^26,27^ Interactions of these three peptides with the experimentally derived TTR tetramer structure as deposited in the Protein Data Bank (PDB) under identifier 1F41 were studied comparing their binding sites with that of thyroxine, the retinol binding protein, and the tetramer-stabilizing tafamidis. Our goal is not only to test whether these small viral protein fragments do de-stabilize the TTR tetramer, but also to identify whether the underlying mechanism exploits weaknesses in the tetramer structure that also are seen in familiar (i.e., mutation-caused) forms of TTR amyloidosis. We find that the viral protein fragments do decrease the stability of the TTR tetramer, potentially easing the way to aggregation and amyloidosis, not by reducing the dimer-dimer interactions but by interfering with transient hydrogen bonding at the monomer-monomer interfaces.

## Materials and Methods

### System preparation

This study employed molecular dynamics to investigate how the stability of transthyretin tetramer complex is altered in the presence of three SARS-CoV-2 derived viral protein fragments. Two viral fragments, ^194^FKNIDGYFKI^203^ (FI10) and ^1058^HGVVFLHVTYV^1068^ (HV11), were extracted from SARS-CoV-2 Spike protein model (PDB ID 6VVX), while the ^55^SFYVYSRVK^63^ (SK9) fragment was obtained from the envelope protein model deposited in the MOLSSI COVID-19 hub.^28^ Licorice representations of the three fragments are shown in **Figure 1a-1c**.

**Figure 1.**
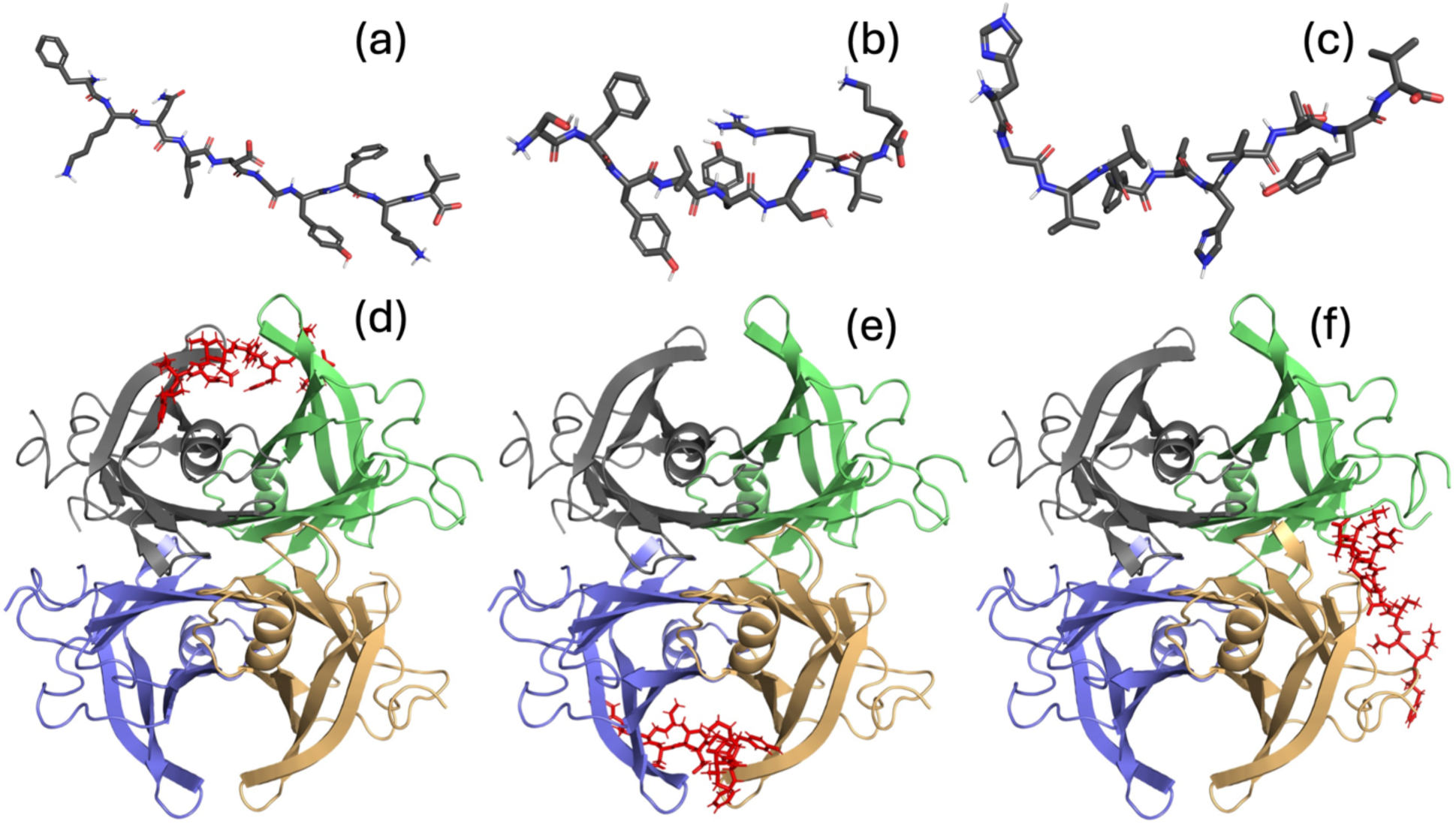
Licorice representation of SARS-CoV-2 derived viral fragments, (a) FI10 (b) SK9, and (c) HV11. Oxygen, Nitrogen, Carbon, and polar hydrogen atoms are drawn in red, blue, grey, and white, respectively. As an example, for the start conformations of our simulations, we show in (d) such with FI10 and TTR binding at the 1-2 monomer interface, in (e) with the binding at the 3-4 interface, and in (f) at the dimer interface. Here, FI10 is drawn in red and the TTR chains are colored in grey (1), lime (2), blue (3), and light orange (4).

As TTR fibril model we considered a wild type TTR structure, resolved by X-ray diffraction with a resolution of 1.3 Å and deposited in the Protein Data Bank under identifier 1F41. The unresolved first nine and the last three residues in each chain were added by us as loops using the MODELLER 10.7^29^ software as implemented within UCSF Chimera.^30^ The resulting structure serves as the start point for our TTR control simulations.

In a next step, we sampled potential binding modes between viral fragments and the TTR tetramer using the HADDOCK 2.4^31,32^ web server while maintaining a 1:1 ratio between tetramer and the viral fragments (one fragment per tetramer), docking viral fragments to either the 1-2 monomer-monomer interface, the dimer-dimer interface, or by assigning randomly active residues spread over the entire TTR surface. Interestingly, in all of the 10 random docking calculations, the viral fragments attached preferentially to the 3-4 monomer interface. Comparing cluster sizes and docking scores, we finally selected three TTR-viral fragment complexes. In the first one, the viral fragments bind to the 1-2 monomer interface, in the second to the 3-4 monomer interface, and in the third complex to the dimer-dimer interface. These three TTR-viral complexes served as the start conformations for our main simulations, and are shown in **Figure 1d-1f** for the example of FI10. Note that the positions of viral protein fragments are not fixed, and that they can move along the tetramer or even detach.

### General simulation protocol

Our simulations rely on the GROMACS 2022 package,^33^ with the CHARMM36m all-atom force field^34^ and the TIP3P explicit water model^35^ used to describe inter-molecular and intra-molecular interactions. The pdb2gmx module in the GROMACS suite was used to add hydrogens.^33^ The resulting configurations were centered in a cubic box with periodic boundary conditions, with at least 15 Å between the solute and edge of the box. The box was then filled with water molecules and Na^+^ and Cl^−^ ions at the physiological ion concentration of 150 mM NaCl. The total number of water molecules and atoms of our systems are listed in the **Table 1**. Each system was equilibrated by the steepest descent algorithm for up to 50000 steps, followed by a 200 ps molecular dynamics simulation at constant volume and 310 K, and another at 310 K and constant pressure (1 bar), constraining heavy atoms positions with a force constant of 1000 kJmol^−1^nm^−2^.

**Table 1.** Simulated Systems.

| System Description |  | Atoms | Water molecules | Independent trajectories | Simulation length (ns) | Total Sampling time (ns) |
| --- | --- | --- | --- | --- | --- | --- |
| Control |  | 103101 | 31741 | 3 | 500 | 1500 |
| FI10 | At 1-2 Interface | 102137 | 31361 | 1 | 500 | 1500 |
|  | At 3-4 Interface | 103004 | 31650 | 1 | 500 |  |
|  | At dimer Interface | 105760 | 32566 | 1 | 500 |  |
| SK9 | At 1-2 Interface | 102638 | 31533 | 1 | 500 | 1500 |
|  | At 3-4 Interface | 102677 | 31546 | 1 | 500 |  |
|  | At dimer Interface | 102398 | 31453 | 1 | 500 |  |
| HV11 | At 1-2 Interface | 102537 | 31493 | 1 | 500 | 1500 |
|  | At 3-4 Interface | 102804 | 31582 | 1 | 500 |  |
|  | At dimer Interface | 103466 | 31802 | 1 | 500 |  |

Following equilibration we performed independent trajectories for each TTR-tetramer systems covering 500 ns at a constant temperature (310 K) and constant pressure (1 bar). The V-rescale thermostat^36^ with a 0.1 ps coupling constant was used to control the temperature during the simulation, while the pressure was handled using the Parrinello-Rahman barostat^37^ with 2 ps relaxation time. Rigidity of the water molecules was maintained by the SETTLE algorithm,^38^ and the LINCS algorithm^39^ was used to restrain protein bonds while keeping hydrogen atoms at the equilibrium distances. This allows us to integrate the equations of motion with a 2 fs time step. To account for the periodicity of the simulation box, we calculated long-range electrostatic interactions by the particle-mesh Ewald (PME) method^40^ with 12 Å real space cutoff and 1.6 Å Fourier grid spacing. The short-range Van der Waals interactions are calculated with the force-switch method where the interactions ae truncated at 12 Å with smoothing starting at 10 Å. Simulation details are also listed in **Table 1**, and the start and final configurations of our trajectories shown in **supplementary material SF1**. The corresponding atomic coordinates are provided in the Protein Data Bank (PDB) format in separated files as **supplementary material.**

### Trajectory Analysis

We visualize our trajectories and conformations using VMD^41^, PyMol^42^, and UCSF Chimera^30^. GROMACS tools were used for the calculation of root mean square deviation (RMSD), root mean square fluctuation (RMSF), solvent accessible surface area (SASA), radius of gyration (Rg), distance between center of masses, and hydrogen bonds. As shown in the results section, the RMSD of the protein backbone has reached a plateau after 300 ns. Therefore, only the last 200 ns were considered for the analysis of our trajectories. The solvent accessible surface area was calculated using a 1.4 Å spherical probe. Hydrogen bond analysis was performed using a 3.5 Å donor-acceptor distance cutoff and an angle cutoff of 30°. We used the MDAnalysis python library^43^ for the calculation of contacts between chains, secondary structure, and residue-wise binding probability. A distance cutoff of 4.5 Å between heavy atoms was used for the calculation of contacts. Secondary structure of the protein was determined using the Dictionary of the Secondary Structure of Proteins (DSSP) algorithm.^44^ Binding probabilities were calculated according to the equation 1 using only the last 200 ns of the trajectories. The NumPy library^45^ was used for data averaging, standard deviation calculation, and other mathematical calculations, while Matplotlib library^46^ was used to plot data.

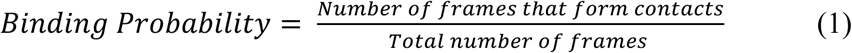

## Results and Discussion

In order to study the effect of SARS-CoV-2 protein fragments on the earliest stages of TTR amyloid formation, we compare all-atom molecular dynamic simulations of the TTR tetramer in presence and absence of three viral protein fragments. Visual inspection of the trajectories and the final conformations (see **supplementary material SF1**) show little changes in the overall structure, but also reveals that the protein fragments can disengage and re-attach. For instance, though FI10 is initially bound at the dimer interface (between chain 2 and 4) in one of our simulations, it detached from the TTR tetramer after about 315 ns, and re-attached to chain 2 at around 425 ns. Defining a viral peptide as bound to the TTR tetramer if at least half of its residues are in contact with TTR chains and using the approach described by Bellaiche and Best^47^, we find as free energy of binding to the tetramer for FI10 −17.9(1) kJ/mol, for SK9 −19.3(1) kJ/mol, and for HV11 −29.4(1) kJ/mol. The stronger binding of HV11 to the TTR chains can be also seen in the binding probabilities of the individual residues to TTR chains in **Figure 2a-2c**. Note the difference between the rather even distributions to the corresponding probability distributions of TTR binding sites in **Figure 2d-2f** where we see a clear clustering that depends on the specific fragments.

**Figure 2.**
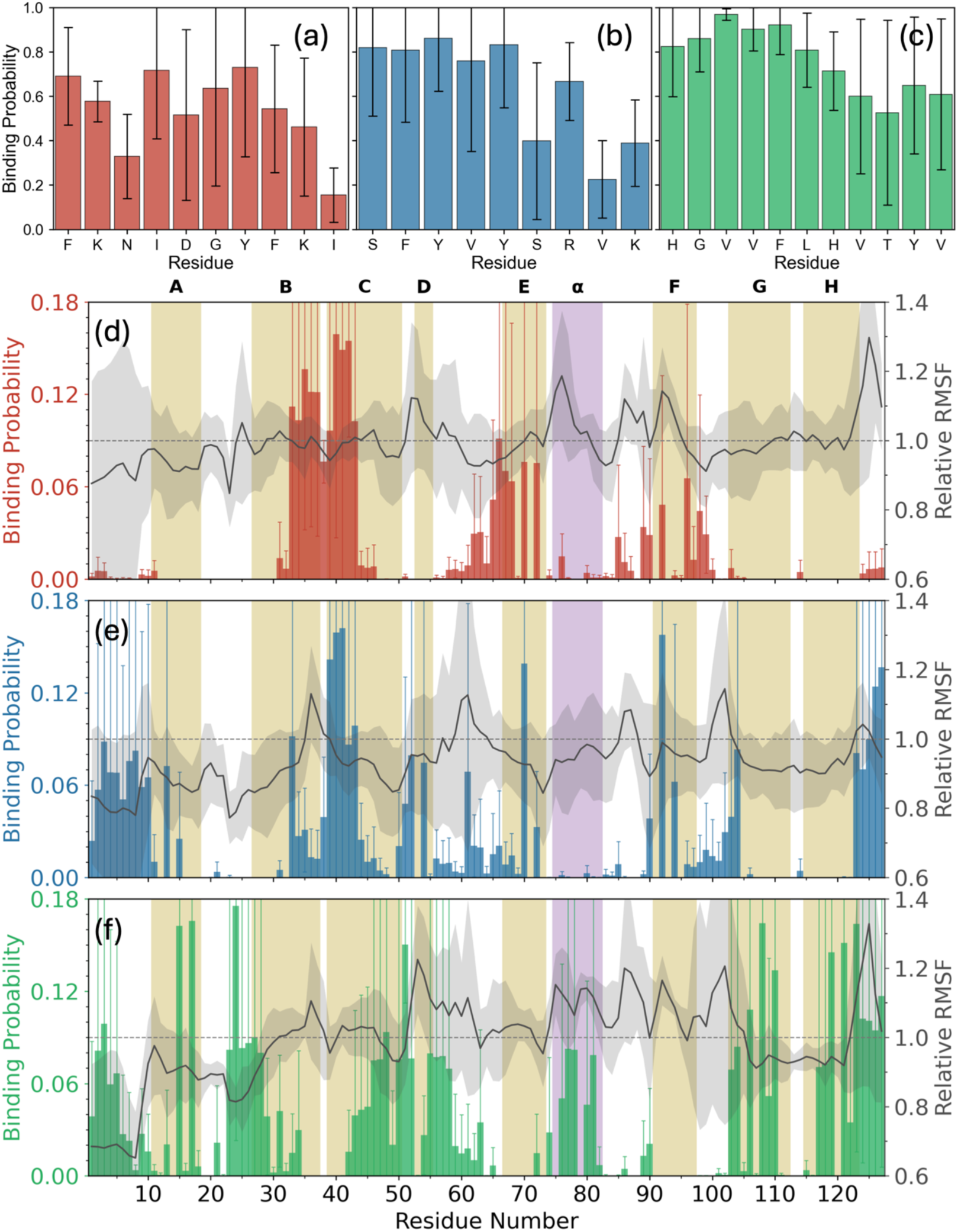
Average binding probability of TTR chains with residues of (a) FI10 (b) SK9 and (c) HV11. Corresponding binding probability of TTR chain residues with (d) FI10, (e) SK9, and (f) HV11 protein fragments. Relative RMSF is shown in black. TTR segments with defined secondary structure are shaded in gold (strands) and purple (helix). Probabilities are calculated over last 200 ns.

The binding probabilities of the TTR binding sites are correlated with regions rich in hydrophobic residues and the strands B (N27-A37) and C (D39-S50), and to a lesser degree F (A91-A97), G (R103-S112) and H (S115-T123). Binding to the *a*-helix (T75-L82) is only seen for HV11, and here only in chain 1 where this binding stabilizes the helix. However, we do not see a clear correlation with the residue-wise root mean square fluctuations (RMSF), a measure for the local flexibility of the TTR chains. The relative RMSF, i.e., the ratio of RMSF measured in presence of a viral fragment divided by the RMSF measured in the control, is also shown in **Figure 2d-2f**, and appears to be not correlated with the binding probabilities. Absolute RMSF values for the control and the three complexes are shown in **supplementary material SF2a -SF2c**. Comparing the clusters of binding sites with the ones of the retinol binding protein RBP and thyroxine, the main binding partners of TTR, or that of tafamidis, the drug used to stabilize the TTR tetramer, all three listed in **supplementary material ST1,** we do not see any competition that would affect the function of TTR.

The changes in the tetramer structures over the course of the trajectories are quantified by the root-mean-square deviations (RMSD) to the resolved crystal structure as function of time shown in **supplemental material SF3a-SF3c**, where we find little differences caused by presence of the protein fragments. In all cases the RMSD values approach a plateau after 300 ns or earlier, and the general behavior stays the same when the RMSD is calculated separately for each chain. We use therefore for calculating equilibrium properties only the last 200 ns of each trajectory.

We list in **Table 2** for a number of quantities the respective values at start together with the average over the last 200 ns of all three trajectories. As a rule, the values do not change much over the course of the simulation, and the effect of the viral protein fragments is small, leading to marginal higher values for Radius of Gyration (Rg) and the solvent accessible surface area (SASA), resulting mostly from hydrophobic residues, indicating slightly less compact structures in presence of the three fragments. We remark that the larger Rg values in presence of the viral peptides result from dimer 1, while the opposite behavior is seen for dimer 2, consistent with larger interface distances between chain 1 and 2, while the values for the 3-4 monomer interface, or for the interface between the two dimers, change little. Correspondingly, we find in presence of the viral protein fragments no reduction in the number of contacts between the two dimers at the dimer interface, but a reduced number of contacts between chain 1 and 2, see **Table 3**. For the contacts between chain 3 and 4 we see the reduction only for FI10. The picture does not change if we consider only native contacts, that is contacts seen already in the resolved crystal structure, and that we do not see significant changes in the number of intra-strand contacts. Hence, the viral protein fragments neither change the geometry of the individual chains nor do they interfere with the stability of the dimer-dimer interface, but they do affect the binding of the monomers 1 and 2, and in the case of FI10 also the binding of chains 3 and 4.

**Table 2.** Radius of gyration (Rg), interfacial distances, and solvent accessible surface area (SASA) measured for the TTR tetramer as averages over the final 200 ns of three independent trajectories.

|  | Control |  | FI10 |  | SK9 |  | HV11 |  |
| --- | --- | --- | --- | --- | --- | --- | --- | --- |
|  | Initial | Last 200 ns | Initial | Last 200 ns | Initial | Last 200 ns | Initial | Last 200 ns |
| Radius of gyration (Å) | 21.8 | 22.2 (1) | 21.8 | 22.3 | 21.8 | 22.3 (1) | 21.8 | 22.3 |
| 1-2 monomer Interface Distance (Å) | 7.9 | 8.1 (4) | 7.9 | 8.6 (1) | 7.9 | 8.5 (1) | 7.9 (1) | 8.6 (3) |
| 3-4 monomer Interface Distance (Å) | 7.9 (1) | 8.3 (4) | 7.9 | 8.7 (2) | 7.8 (1) | 8.5 (5) | 7.8 (1) | 8.4 (4) |
| Dimer Interface Distance (Å) | 6.1 | 6.2 (1) | 6.1 | 6.3 (1) | 6.1 | 6.3 (2) | 6.1 (1) | 6.2 (1) |
| Total SASA (nm<sup>2</sup>) | 196 (1) | 208 (4) | 192 | 212 (2) | 192 (1) | 210 (3) | 192 | 212 (3) |
| Hydrophobic SASA (nm<sup>2</sup>) | 88 (0) | 96 (2) | 89 | 97 (2) | 89 (1) | 96 (2) | 89 | 98 (2) |
| Hydrophilic SASA (nm<sup>2</sup>) | 109 (1) | 113 (3) | 103 (1) | 115 | 102 (1) | 114 (2) | 103 | 114 (1) |

This is a surprising result as commonly is assumed that the dimer-dimer interface is more crucial for the stability of the tetramer than the monomer-monomer interfaces in this dimer of dimers.^11,48^ TTR dissociation at the monomer-monomer interface is considered unlikely because of the strong hydrogen bond networks between H and F strands of chain 1 and chain 2, and between chain 3 and chain 4.^48,49,50^ On the other hand, the two dimers assemble through much weaker hydrophobic interactions between the AB loop (residues A19-I26 connecting strands A and B) in chain 1 with the GH loop (residues P113-Y114 connecting strands G and H) in chain 4, and similarly between chain 2 and 3.^11,48,51,52^ Note that both thyroxine and the drug tafamidis bind to the hydrophobic pocket located at this weak dimer-dimer interface stabilizing the interface.

Hence, despite the dimer interface being considered the weak point in the TTR tetramer, our above contact analysis indicates that the viral protein fragments destabilize not this interface but the monomer-monomer interface. While this is a surprising result, we remark that the monomer-monomer interface is also vulnerable to mutations such as L55P or V30M, which potentially affect the equilibrium dynamics of TTR tetramer.^51,53,54^ Note also that we observed a larger effect of the viral peptides on the 1-2 monomer interface than on the 3-4 monomer interface, indicating that despite the 2-fold symmetry of the tetramer, the constituting dimers may have different dynamics as was also suggested in earlier work.^54^

Consistent with our hypothesis that the viral protein fragments destabilize the tetramer by weakening the monomer-monomer, we find in **Table 3** follows that the viral peptides interfere with the hydrogen bonds connecting strands F and H on chain 1 and 2, and in case of FI10 also chain 3 and 4. Note that for HV11 we find at start two more hydrogen bonds than in the control or in presence of SK9 or FI10. This is an artefact of how we identify hydrogen bonds and the difference disappears if the angle criteria is slightly relaxed.

**Table 3.** Hydrogen bonds (H-bonds), contacts, and native contacts measured for the TTR tetramer as averages over the final 200 ns of three independent trajectories.

|  | Control |  | FI10 |  | SK9 |  | HV11 |  |
| --- | --- | --- | --- | --- | --- | --- | --- | --- |
|  | Initial | Last 200 ns | Initial | Last 200 ns | Initial | Last 200 ns | Initial | Last 200 ns |
| H-bonds 1-2 Monomer Interface | 12.7 (6) | 14.6 (2.4) | 12.7 (6) | 11.4 (7) | 11 (1) | 11.9 (6) | 14 (2) | 11.6 (5) |
| H-bonds 3-4 Monomer Interface | 11.7 (6) | 13.0 (2.4) | 11.3 (2.1) | 11.1 (9) | 13.3 (6) | 12.8 (2.9) | 14 (3) | 12.2 (1.2) |
| Contacts 1-2 monomer interface | 56.7 (6) | 53 (7) | 55(2) | 44 (3) | 55.7 (6) | 46 (2) | 56 (2) | 46(2) |
| Native contacts 1-2 monomer interface | 56 (1) | 44.5 (2.7) | 54 (2) | 41.4 (2.5) | 54.3 (6) | 42.2 (1.2) | 54.3 (2.1) | 41 (4) |
| Contacts 3-4 monomer interface | 56 (2) | 49(6) | 55.3 (2.1) | 43.4 (3.2) | 58.3 (1.2) | 48 (8) | 55(1) | 47 (5) |
| Native contacts 3-4 monomer interface | 55 (2) | 42.3 (3.1) | 55 (2) | 40.6 (3.7) | 56.3 (6) | 42.5 (3.6) | 55(1) | 43(3) |
| Contacts dimer interface | 75 (1) | 85.9 (2.5) | 73(2) | 86.7 (2.0) | 72.7 (2.3) | 84.3 (2.3) | 74(4) | 85.9 (1.0) |
| Native contacts dimer interface | 71.3 (6) | 66.5 (0.9) | 69.3 (6) | 65.6 (2.7) | 69.3 (1.5) | 64.9 (3.6) | 70.7 (1.5) | 66.5 (9) |
| Intra-strand contacts | 274(1) | 260 (3) | 275 (3) | 257.7 (1.6) | 274 (1) | 260(4) | 274(3) | 255 (1) |
| Native<br>intra-strand<br>contacts | 267 (1) | 234(3) | 265 (3) | 232.1<br>(2.7) | 264(1) | 233 (5) | 265(4) | 228 (1) |

Note that the standard deviation in the number of hydrogen bonds between chain 1 and 2 in presence of the viral peptides is smaller than in the control. A more detailed analysis of the control trajectories (i.e, in absence of any interacting peptides) shows indeed a structural diversity of the tetramer configurations over the course of the trajectories. For instance, in two of the three trajectories we find similar distributions of Rg while the third trajectory leads to a distribution that is shifted to larger values, see **Figure 3a**. This shift is also seen when the TTR tetramer is interacting with the viral protein fragment, see **Figure 3b-3d**. We remark that these differences in the Rg distribution are correlated with similar differences in the distribution of interfacial distance, hydrogen bonds, contacts, native contacts, and interfacial SASA (data not shown).

**Figure 3.**
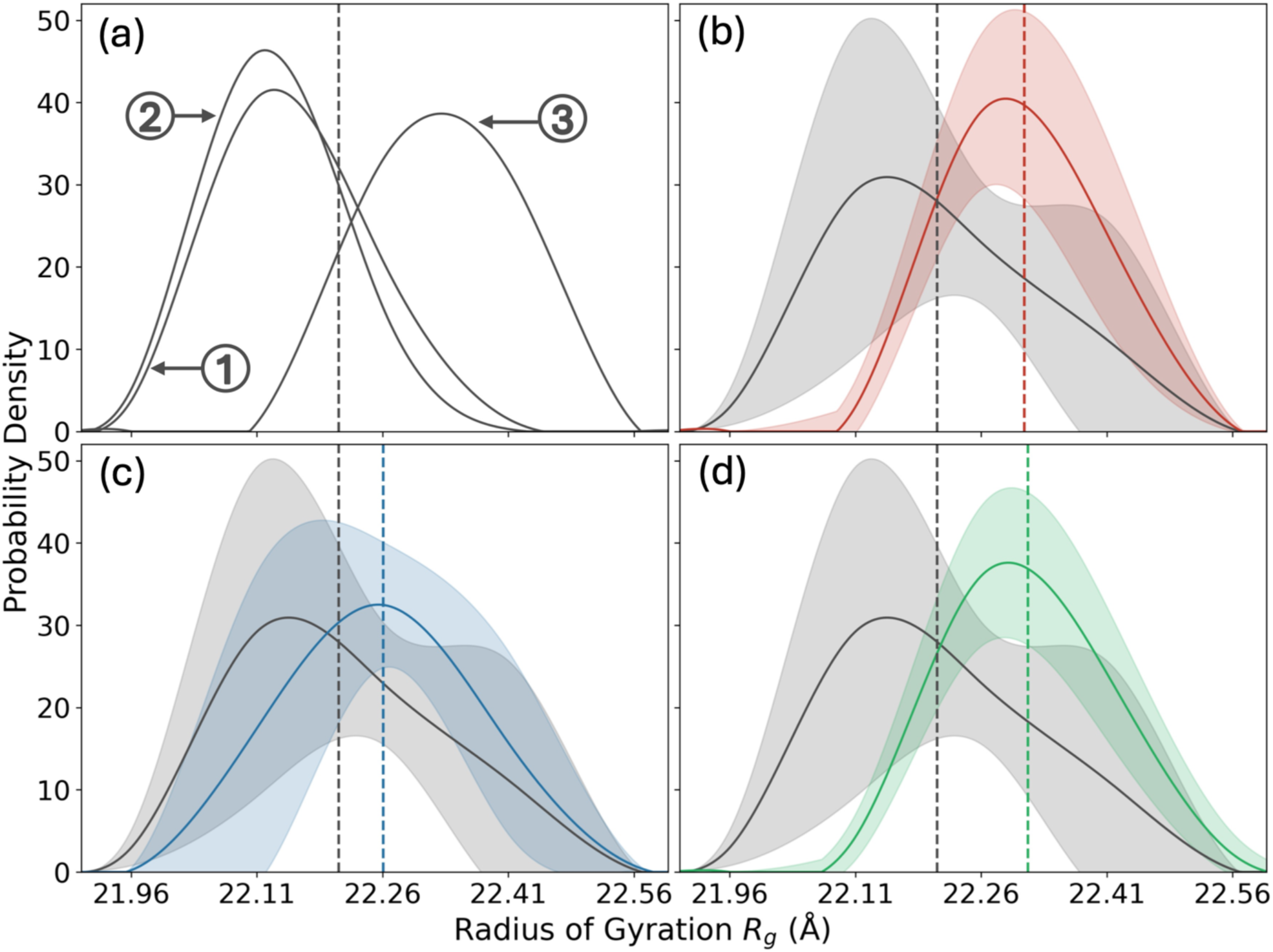
Probability density of (a) radius of gyration of individual trajectories of the control simulations. Average distributions of the control simulations are compared in (b)-(d) with the corresponding distributions of simulations where viral protein fragments interact with the TTR tetramer, FI10 in (b), SK9 in (c), and HV11 in (d). The average radius of gyration calculated over last 200 ns of the control, FI10, SK9, and HV11 simulations is indicated by black, red, blue, and green verticle lines, respectively.

For the control simulations, the shift in the Rg distributions is related to a loss of hydrogen bonds connecting the F strand on chain 1 and 2 (or chain 3 and 4). This can be seen in **Figure 4** where we show in **Figure 4a**-**4c** the frequency of hydrogen bonds between residues at the 1-2 monomer interface in the three control trajectories, and in **Figure 4d** the difference between these frequencies measured in the third trajectory and the average of the corresponding frequencies measured in the first two trajectories. Corresponding contact maps for the 3-4 interface are shown in **supplementary material SF4**. Differences in the hydrogen bond frequency between third and the first two trajectories are most pronounced for residue pairs E89-T96, H90-V94, E92-E92, V94-H90 and T96-E89, and to a lesser degree between residues H88-T118 and T118-H88, see **Table 4**.

**Figure 4.**
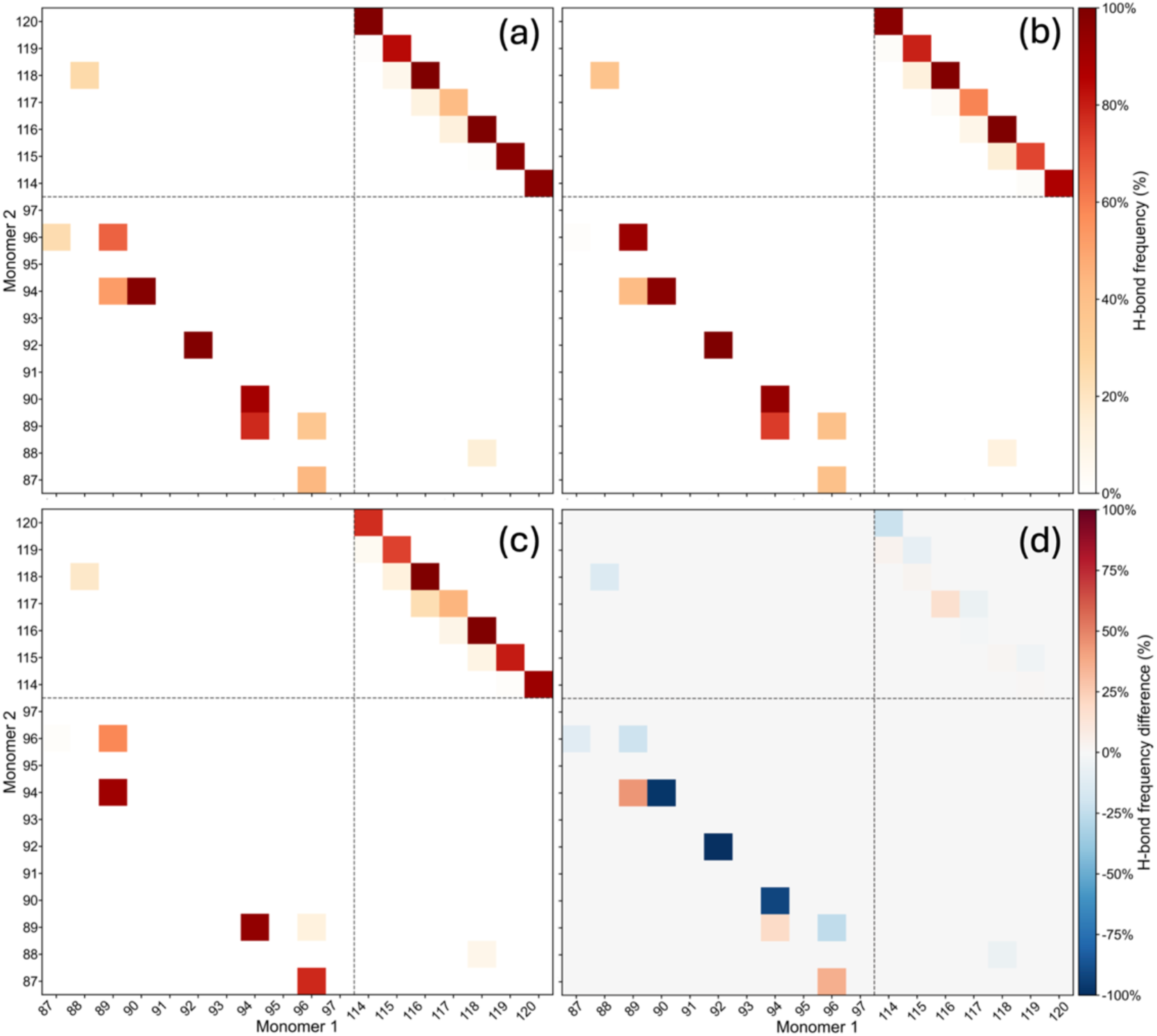
Hydrogen bond frequency at the 1-2 interfaces as measured in the control simulations for (a) trajectory 1 (b) trajectory 2, and (c) trajectory 3. In (d) we show the difference in hydrogen bond frequency between trajectory 3 and the average of trajectory 1 and 2.

**Table 4.** Hydrogen bond frequencies at the 1-2 and the 3-4 monomer-monomer interface as measured in the three control simulations C1, C2 and C3. We also list the difference between values measured in run 3 and the average of run 1 and run 2.

| Residue Pair | Hydrogen bond frequency % monomer1-monomer2 Interface |  |  |  | Hydrogen bond frequency % monomer3-monomer4 Interface |  |  |  |
| --- | --- | --- | --- | --- | --- | --- | --- | --- |
| | C1 | C2 | C3 | $C3-(C2+C1)/2$ | C1 | C2 | C3 | $C3-(C1+C2)/2$ |
| H88-T118 | 26 | 38 | 18 | -13 | 8 | 21 | 18 | 3 |
| E89-T96 | 66 | 92 | 58 | -21 | 29 | 28 | 72 | 44 |
| H90-V94 | 98 | 96 | 0 | -97 | 0 | 92 | 54 | 8 |
| E92-E92 | 99 | 100 | 0 | -99 | 0 | 99 | 0 | -50 |
| V94-H90 | 89 | 94 | 0 | -91 | 0 | 99 | 0 | -50 |
| T96-E89 | 36 | 39 | 12 | -25 | 8 | 59 | 11 | -22 |
| T118-H88 | 15 | 12 | 8 | -6 | 0 | 20 | 13 | 3 |

Comparing in the control simulations the structural variation with changes in frequency of the hydrogen bonds, it appears that the hydrogen bonds listed in **Table 4** are important for the stability of the tetramer. This assumption is consistent with a recent joint neutron and X-ray crystallographic analysis^55^ that found stabilization of the monomer interface through direct and water mediated hydrogen bonds formed between F strands on chain 1 and 2 (or chain 3 and 4), including the residue pairs H90-V94, E92-E92, V94-H90 discussed above. The monomer interface is also stabilized by hydrogen bonds formed between H strands of the two interacting chains including one between residues T119-Y114, S115-T119, S115-T118, Y116-T118, T118-S115, T118-Y116, and T119-S115; and by a water mediated hydrogen bonds formed between F and H strands of opposite chains such as between residues H88 -T118.^55^

The importance of the above discussed hydrogen bonds is supported by comparing pathogenic mutations with wild type TTR. For instance, a NMR study has demonstrated that V30M, the most common pathogenic mutation, and L55P, the most aggressive mutation, induce a significant ^15^N, ^1^H_N_, and ^13^C*α* chemical shift perturbation in E89, V93, S115, S117, and T118 residues, indicating structural changes that could potentially interrupt the hydrogen bonding at monomer interface.^51^ Another study has shown that the frequency of T96-F87 hydrogen bond in dimer 1, and, V90-H94, E92-E92, H94-V90, and T96-F87 hydrogen bonds in dimer 2 in the L55P mutant is lower than in wild type TTR.^54^ Frequency of hydrogen bonds H90-V94, E92-E92, and V94-H90, T96-F87 in dimer 1, and V90-H94 and E92-E92 in dimer 2 are also reduced in the V30M variant.^54^

Our comparison of the control trajectories suggests that interference with the above hydrogen bonds could be a mechanism by which the viral protein fragments may destabilize the TTR tetramer. This assumption is supported by the observation that the frequencies of these hydrogen bonds are also reduced in the presence of the viral protein fragments in **Figure 5a-5c** and **supplementary material SF5**. We find that, for instance, in dimer 1 (chain 1 and 2), the hydrogen bond frequency between residue V94 in monomer 1 and residue H90 in monomer 2 (V94-H90), drops from 61% in the control simulations to almost 0% in all viral fragment simulations. In dimer 2 (chain 3 and 4), we find a similar drop in the frequency from 33% to values between 0% (FI10) and 7% (HV11). However, there was almost no drop for SK9 where we observed 31% frequency for the V94-H90 hydrogen bond. While the hydrogen bonding between H strands is not affected by the presence of viral protein fragments, the hydrogen bond formed between F and H strands that involved residue H88 of chain 1 and T118 of chain 2 (H88-T118) has dropped from 27% in control simulations to 8% in FI10, 14% in HV11 simulations and to 24% in SK9 simulations. The frequencies of these hydrogen bond pairs as measured in presence of the viral peptides are listed in **Table 5**. Note that we do not see the expected reduction in frequency for the pair T96-E89 but instead one for T96-F87.

**Figure 5.**
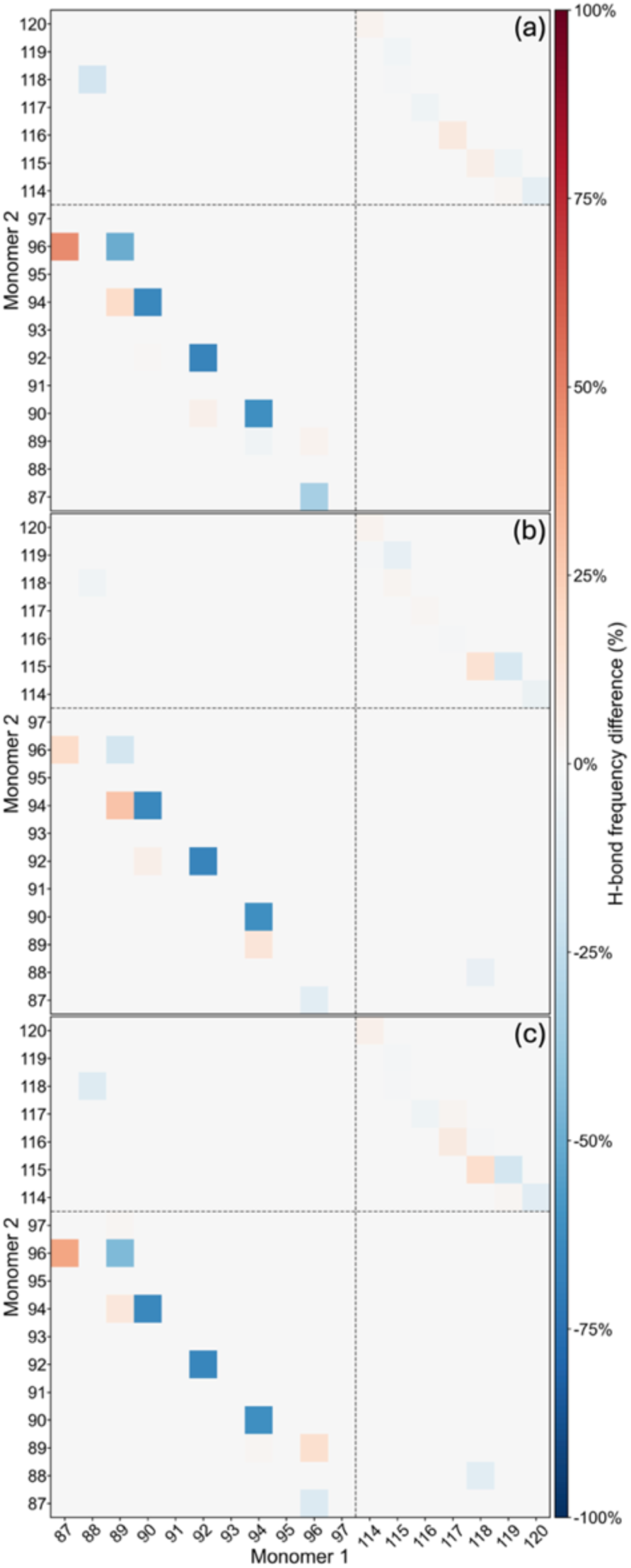
Difference in hydrogen bond frequency at 1-2 monomer interface between simulations where (a) FI10, (b) SK9, and (c) HV11 are present, and the control simulations.

**Table 5.** Frequencies of the discussed hydrogen bonds between F-strands at the monomer-monomer interfaces.

| Residue Pair | Hydrogen bond frequency % monomer1-monomer2 Interface |  |  |  | Hydrogen bond frequency % monomer3-monomer4 Interface |  |  |  |
| --- | --- | --- | --- | --- | --- | --- | --- | --- |
|  | Control | FI10 | SK9 | HV11 | Control | FI10 | SK9 | HV11 |
| H88-T118 | 27 (10) | 8 (14) | 24 (14) | 14 (25) | 16 (7) | 1 (2) | 26 (31) | 19 (20) |
| E89-T96 | 72 (18) | 22 (6) | 54 (20) | 28 (28) | 43 (25) | 28 (27) | 42 (37) | 35 (23) |
| H90-V94 | 65 (56) | 0 (0) | 0 (0) | 0 (0) | 49 (46) | 0 (0) | 11 (19) | 7 (12) |
| E92-E92 | 66 (57) | 0 (0) | 0 (1) | 1 (1) | 33 (57) | 0 (0) | 33 (58) | 8 (15) |
| V94-H90 | 61 (53) | 0 (0) | 0 (0) | 0 (0) | 33 (57) | 0 (0) | 31 (54) | 7 (12) |
| T96-F87 | 54 (21) | 21 (34) | 43 (46) | 40 (42) | 37 (47) | 58 (47) | 20 (16) | 29 (50) |
| T96-E89 | 29 (15) | 33 (29) | 30 (41) | 45 (34) | 26 (29) | 23 (32) | 32 (10) | 38 (32) |
| T118-H88 | 11 (4) | 11 (15) | 4 (4) | 1 (1) | 11 (10) | 3 (5) | 9 (12) | 22 (35) |
| **Conditional hydrogen bond frequencies when the peptides bind to residues 27-50 (strands B and C)** |  |  |  |  |  |  |  |  |
| H88-T118 | 27 (10) | 8 (14) | 31 (27) | 0 (0) | 16 (7) | 0 (0) | 51 (0) | 0 (0) |
| E89-T96 | 72 (18) | 15 (15) | 56 (30) | 11 (11) | 43 (25) | 0 (0) | 85 (0) | 18 (0) |
| H90-V94 | 65 (56) | 0 (0) | 0 (0) | 0 (0) | 49 (46) | 0 (0) | 16 (0) | 0 (0) |
| E92-E92 | 66 (57) | 0 (0) | 1 (1) | 0 (0) | 33 (57) | 0 (0) | 99 (0) | 0 (0) |
| V94-H90 | 61 (53) | 0 (0) | 0 (0) | 0 (0) | 33 (57) | 0 (0) | 91 (0) | 0 (0) |
| T96-F87 | 54 (21) | 20 (35) | 44 (45) | 43 (61) | 37 (47) | 87 (0) | 13 (0) | 0 (0) |
| T96-E89 | 29 (15) | 46 (39) | 28 (39) | 55 (54) | 26 (29) | 4 (0) | 37 (0) | 35 (0) |
| T118-H88 | 11 (4) | 4 (6) | 4 (4) | 0 (0) | 11 (10) | 0 (0) | 12 (0) | 3 (0) |

By what mechanism does interaction with the viral peptides cause the weakening and loss of these hydrogen bonds between strands F of different chains the monomer-monomer interfaces? While binding only with small probability to strand F, the binding distributions in **Figure 2d-2f** show preferential binding to a segment rich in hydrophobic residues including the strands B (N27-A37) and C (D39-S50), which are part of the CBEF β-sheet. Hence, binding of the viral peptides to the segment of residues N27-S50 could transmit to strand F in a similar way to how the V30M mutation expands slightly the hydrophobic core by moving the CBEF β-sheet away from DAGH β-sheet, and where these perturbations transmit across the CBEF sheet to residues E72, I73 on strand E and to residues E92 and V93 residues in strand F.^51^ In a similar way, the L55P mutation alters not only hydrogen bonding of residues in strand D but the disturbances propagate across both β-sheets.^51^ Our hypothesis is that binding of viral peptides to the B and C strands has the same effect. To test this hypothesis, we have calculated the conditional probabilities for the hydrogen bonds listed in **Table 5** under the condition that a viral peptide binds to residues in the segment N27-S50, and compared these frequencies again with the corresponding frequencies measured in the control. These conditional frequencies are also listed in **Table 5** and show indeed that binding to residues N27-S50 causes a reduction of the hydrogen bonds connecting residues located on the F strands on chains 1 and 2, and to a lesser degree between such on chain 3 and 4, therefore destabilizing hydrogen bonds at the monomer-monomer interface that are already weak or transient in the control.

## Conclusions

Unfolding and aggregation of the TTR chains released after dissociation of the home tetramer is the cause of TTR amyloidosis that especially in elderly patients may damage the heart and other organs. As correlations between TTR amyloidosis and SARS-CoV-2 infections have been observed, ^24^ we have used molecular dynamics simulations to study the effect of three fragments from the spike and envelope proteins of the SARS-CoV-2 virus on the stability of the TTR tetramer, a dimer of dimers. We find little differences betwee the viral protein fragments, but their presence results in less compact structures of the TTR tetramer, with the difference resulting mainly from dimer 1, and here from a separation between the two monomers. This is a surprising result as the monomers in each dimer are stronger bound than the two dimers in the tetramer, and existing drugs are designed to stabilize the dimer-dimer interface. We identify a set of hydrogen bonds between residues on the F-strand on chain 1 and chain 2 (and to a lesser degree chain 3 and chain 4) that already in the control simulations are transient, and where the frequency is reduced in presence of the viral protein fragments. We show that binding of the viral protein fragments to the segment of residues N27-S50, which is rich in hydrophobic residues and includes strands B and C, is correlated with this reduction in hydrogen bonds at the monomer-monomer interface, and pose that the binding introduces a perturbation of the strands that transfers along the β-sheet CBEF weakening these critical interchain hydrogen bonds. While this mechanism for destabilizing the tetramer by the viral fragments is unexpected, it supports previous work pointing out the importance of hydrogen bonding at the monomer-monomer interface^49,50^ and the dynamic asymmetry between dimer 1 and dimer 2.^54^ It also suggests that the location of disease causing mutations (as, for instance, V30M and L55P) and their vicinity are likely points where pathogens could trigger disruption of the TTR tetramer or that potential drugs could stabilize.

## SUPPLEMENTARY MATERIAL

**Supplementary Figures (SF):** start and final tetramer conformations, residue-wise root-mean-square fluctuations over final 200ns for all trajectories, time evolution of root-mean-square deviation to start conformation, hydrogen bond frequencies for the interface between chain 3 and chain 4; **Supplementary Tables (ST):** Binding sites of thyroxine, tafamidis and the retinol binding protein; **README** for a separate folder with atomic coordinates (in PDB format) of initial and final conformations, topology files, and md parameter files of all TTR simulations discussed in the article.

## ACKNOWLEDGMENTS

Our simulations were done using the SCHOONER cluster of the University of Oklahoma, the PETE cluster at Oklahoma State University, and ACCESS resources allocated under grant MCB160005 (National Science Foundation).

## AUTHOR DECLARATIONS

### Author Contributions

**Malinda B. Premathilaka:** Formal analysis (equal); Investigation (equal); Visualization (lead); Writing – original draft (equal); **Ulrich H.E. Hansmann:** Conceptualization (lead); Funding acquisition (lead); Resources (lead); Supervision (lead); Writing – original draft (equal); Writing – review and editing (lead).

## DATA AVAILABILITY

The data supporting the results of this study are available in the Supporting Information and are publicly accessible at https://github.com/ouhansmannlab/ttr_tetramer_viral

