## Supporting tables and figures for "Early Steps in Pathogen-induced TTR-amyloid-formation"

Supplementary material for this article consists of 1 table and 5 figures listed on the following pages, and an additional file (**AB-start-final-coordinates.zip**) containing a compressed folder with the atomic coordinates of the start and final configurations (as ASCII files in the PDB), topology files, and parameter files of all molecular dynamics simulations considered in this study. The folder contains also README files to help identifying the various files.

**Table of content of supplementary material**

**Figures:**

**SF1.** Start configuration and final configuration of TTR tetramer for the control simulations, and in presence of the viral protein fragments FI10,SK9 and HV11. The four TTR chains are colored in grey (1), lime (2), blue (3), and light orange (4). S4

**SF2.** Absolute values of the residue-wise root-mean-square fluctuations (RMSF) calculated over the last 200 ns and averaged over three trajectories, (a) FI10 (red), (b) SK9 (blue), and (c) HV11 (green) simulations. The corresponding values for the control simulations are drawn in black. S5

**SF3.** Time series of the root-mean-square deviation (RMSD) of resolved backbone atoms for simulations where the TTR fibril interacts with viral protein fragment (a) FI10 (red), (b) SK9 (blue), and (c) HV11 (green). The corresponding RMSD for the control simulations is drawn in black. RMSD values are calculated considering the crystal structure as the reference and averaged over three trajectories. S5

**SF4.** Hydrogen bond frequency at the 3-4 interfaces as measured in the control simulations for (a) trajectory 1 (b) trajectory 2, and (c) trajectory 3. In (d) we show the difference in hydrogen bond frequency between trajectory 3 and the average of trajectory 1 and 2. S6

**SF5.** Difference in hydrogen bond frequency at 3-4 monomer interface between simulations where (a) FI10, (b) SK9, and (c) HV11 are present, and the control simulations. S7

**Tables:**

**ST1:** Binding site residues of *t*hyroxine, tafamidis and the retinol binding protein. Both thyroxine and tafamidis share the same binding pocket. S8

**README** for additional files in the compressed folder ttr_coordinates.zip (available as separate file). S9

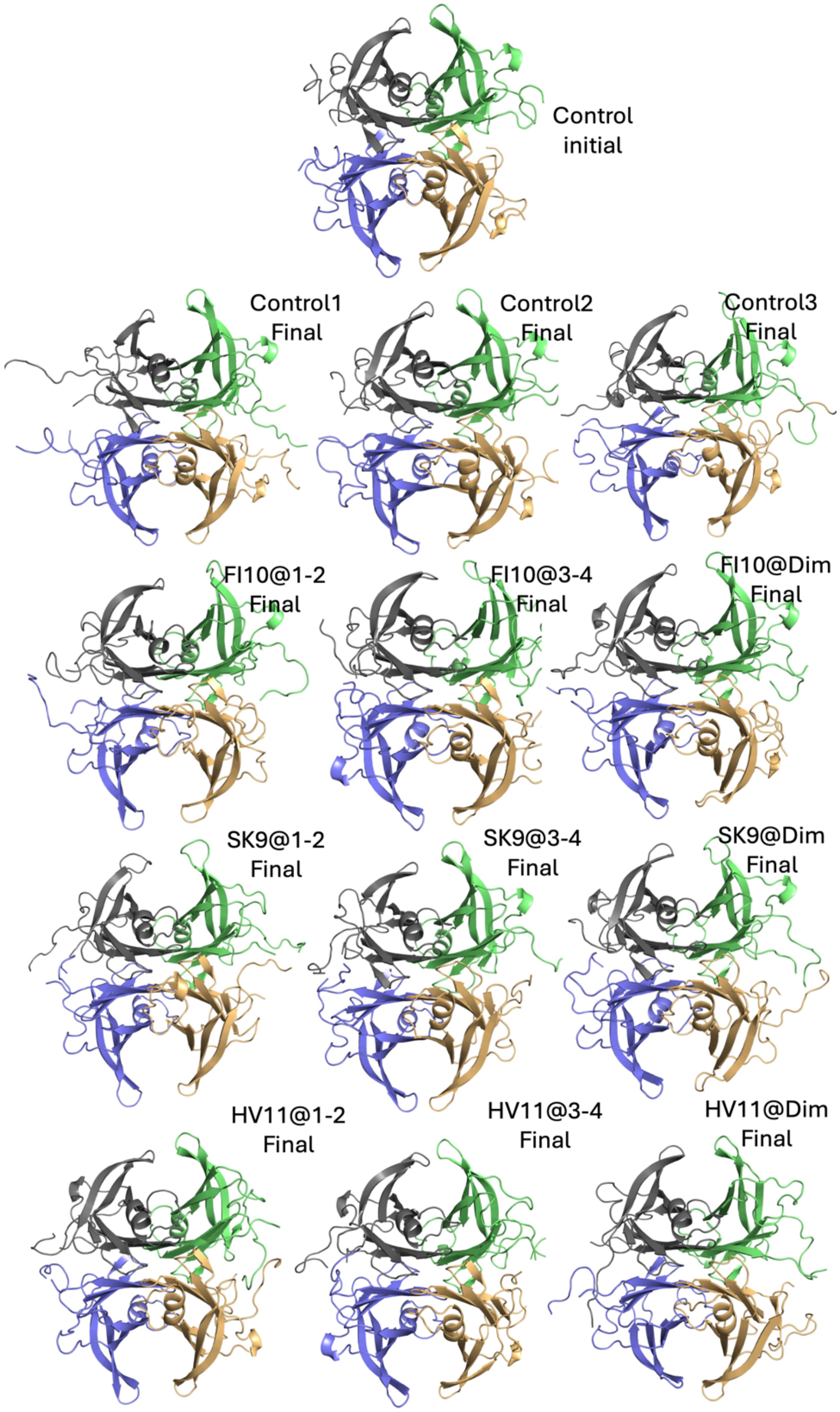


**Figure SF1.** Start configuration and final configuration of TTR tetramer for the control simulations, and in presence of the viral protein fragments FI10,SK9 and HV11. The four TTR chains are colored in grey (1), lime (2), blue (3), and light orange (4).

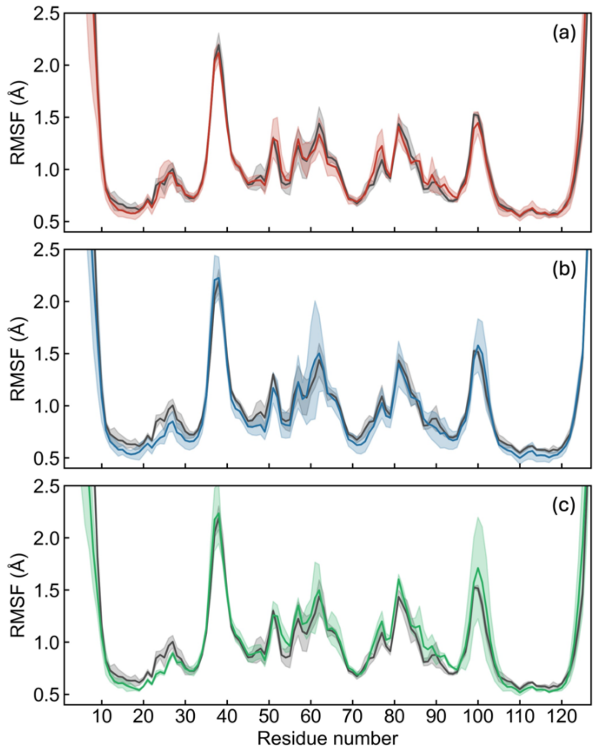


**Figure SF2.** Absolute values of the residue-wise root-mean-square fluctuations (RMSF) calculated over the last 200 ns and averaged over three trajectories, (a) FI10 (red), (b) SK9 (blue), and (c) HV11 (green) simulations. The corresponding values for the control simulations are drawn in black.

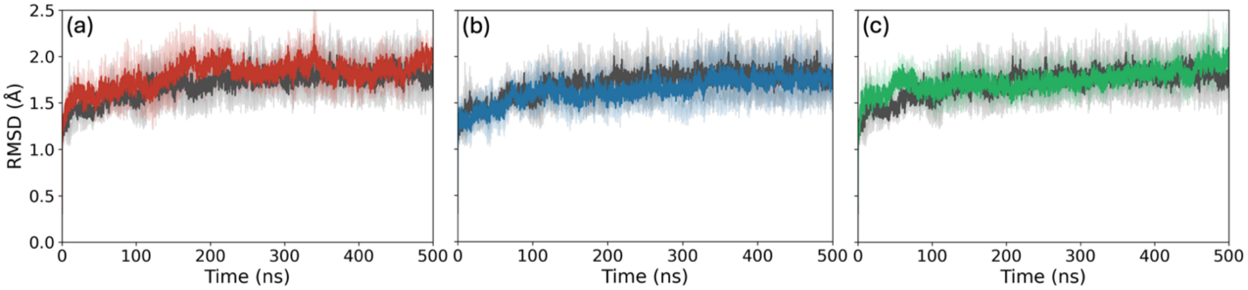


**Figure SF3.** Time series of the root-mean-square deviation (RMSD) of resolved backbone atoms for simulations where the TTR fibril interacts with viral protein fragment (a) FI10 (red), (b) SK9 (blue), and (c) HV11 (green). The corresponding RMSD for the control simulations is drawn in black. RMSD values are calculated considering the crystal structure as the reference and averaged over three trajectories.

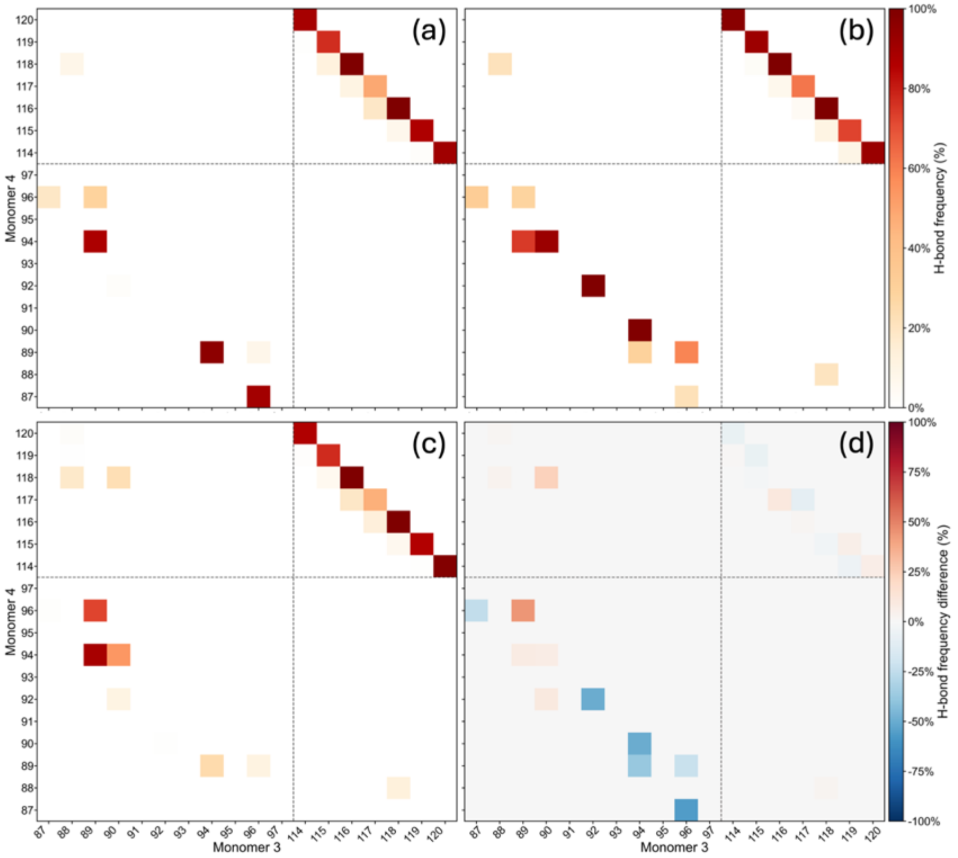


**Figure SF4.** Hydrogen bond frequency at the 3-4 interfaces as measured in the control simulations for (a) trajectory 1 (b) trajectory 2, and (c) trajectory 3. In (d) we show the difference in hydrogen bond frequency between trajectory 3 and the average of trajectory 1 and 2.

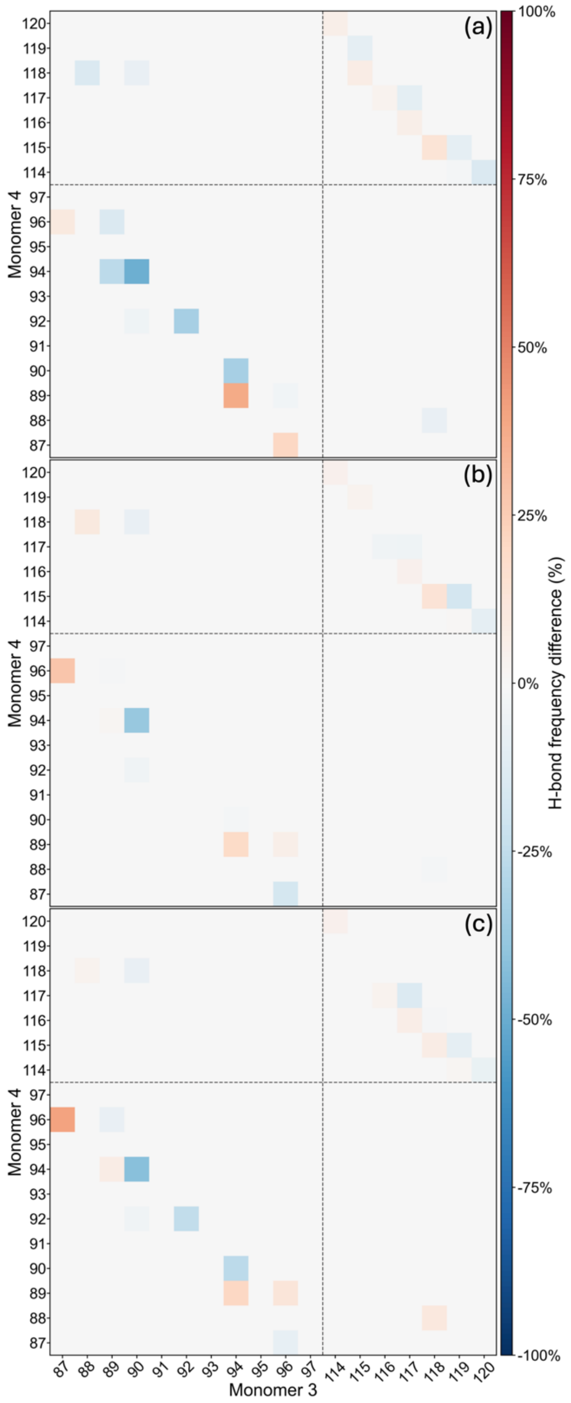


**Figure SF5.** Difference in hydrogen bond frequency at 3-4 monomer interface between simulations where (a) FI10, (b) SK9, and (c) HV11 are present, and the control simulations.

**Table ST1.** Binding site residues of thyroxine, tafamidis and the Retinol Binding Protein. Both thyroxine and tafamidis share the same binding pocket.

| Molecule | Interacting Residues |
|---|---|
| Thyroxine (T4) and Tafamidis | Pocket 1 - M13, K15, T106, A108<br>Pocket 2 – K15, L17, A108, A109, L110,<br>Pocket 3 – A108, A109, L110, S117, T118, T119 |
| Retinol Binding Protein | Chain1 - V20, R21, L82, I84, A81, G83, S85, P86, K76, K80, Y114<br>Chain3 - T96, D99, S100, R103 |

**README for additional files in the compressed folder ttr_coordinates.zip (available as separate file).**

This folder contains (besides this README.txt file) three directories ( (1) coordinates, (2) topology_files, and (3) md_parameter_files) that contain atomic coordinates (in PDB format) of initial and final conformations, topology files, and md parameter files of all ttr simulations discussed in the article.

1. coordinates : This directory contains two sub directories ( (a) initial and (b) final) collecting

    a. initial : contains following files with the coordinates of initial conformations in all simulations.
        i. control_t1_initial.pdb : Initial structure of control simulation trial 1
        ii. control_t2_initial.pdb : Initial structure of control simulation trial 2
        iii. control_t3_initial.pdb : Initial structure of control simulation trial 3
        iv. fi10_dim_initial.pdb : Initial structure of ttr-fi10 dim simulation (fi10 binds to dimer-dimer interface at the beginning)
        v. fi10_mon_initial.pdb : Initial structure of ttr-fi10 mon simulation (fi10 binds to 1-2 monomer interface at the beginning)
        vi. fi10_random_initial.pdb : Initial structure of ttr-fi10 random simulation (fi10 binds to 3-4 monomer interface at the beginning)
        vii. sk9_dim_initial.pdb : Initial structure of ttr-sk9 dim simulation (sk9 binds to dimer-dimer interface at the beginning)
        viii. sk9_mon_initial.pdb : Initial structure of ttr-sk9 mon simulation (sk9 binds to 1-2 monomer interface at the beginning)
        ix. sk9_random_initial.pdb : Initial structure of ttr-sk9 random simulation (sk9 binds to 3-4 monomer interface at the beginning)
        x. hv11_dim_initial.pdb : Initial structure of ttr-hv11 dim simulation (hv11 binds to dimer-dimer interface at the beginning)
        xi. hv11_mon_initial.pdb : Initial structure of ttr-hv11 mon simulation (hv11 binds to 1-2 monomer interface at the beginning)
        xii. hv11_random_initial.pdb : Initial structure of ttr-hv11 random simulation (hv11 binds to 3-4 monomer interface at the beginning)
    b. final : contains following files with the coordinates of final conformations in all simulations.
        i. control_t1_final.pdb : Final structure of control simulation trial 1
        ii. control_t2_final.pdb : Final structure of control simulation trial 2
        iii. control_t3_final.pdb : Final structure of control simulation trial 3
        iv. fi10_dim_final.pdb : Final structure of ttr-fi10 dim simulation (fi10 binds to dimer-dimer interface at the beginning)
        v. fi10_mon_final.pdb : Final structure of ttr-fi10 mon simulation (fi10 binds to 1-2 monomer interface at the beginning)
        vi. fi10_random_final.pdb : Final structure of ttr-fi10 random simulation (fi10 binds to 3-4 monomer interface at the beginning)
        vii. sk9_dim_final.pdb : Final structure of ttr-sk9 dim simulation (sk9 binds to dimer-dimer interface at the beginning)

viii. sk9_mon_final.pdb : Final structure of ttr-sk9 mon simulation (sk9 binds to 1-2 monomer interface at the beginning)
ix. sk9_random_final.pdb : Final structure of ttr-sk9 random simulation (sk9 binds to 3-4 monomer interface at the beginning)
x. hv11_dim_final.pdb : Final structure of ttr-hv11 dim simulation (hv11 binds to dimer-dimer interface at the beginning)
xi. hv11_mon_final.pdb : Final structure of ttr-hv11 mon simulation (hv11 binds to 1-2 monomer interface at the beginning)
xii. hv11_random_final.pdb : Final structure of ttr-hv11 random simulation (hv11 binds to 3-4 monomer interface at the beginning)

2. topology_files : Contains topology files for each simulations. Files are organized according to the simulations as described below. The following subdirectories are available in this directory.

   a. control/trial_1
   b. control/trial_2
   c. control/trial_3
   d. fi10/dim
   e. fi10/mon
   f. fi10/random
   g. sk9/dim
   h. sk9/mon
   i. sk9/random
   j. hv11/dim
   k. hv11/mon
   l. hv11/random

   In each of the above subdirectories, following files are available.

   - topol.top : Topology file of the system
   - topol_Protein_chain_A.itp : Topology file of chain 1 of ttr tetramer
   - topol_Protein_chain_B.itp : Topology file of chain 2 of ttr tetramer
   - topol_Protein_chain_C.itp : Topology file of chain 3 of ttr tetramer
   - topol_Protein_chain_D.itp : Topology file of chain 4 of ttr tetramer
   - posre_Protein_chain_A.itp : Position restraint file of chain 1 of ttr tetramer
   - posre_Protein_chain_B.itp : Position restraint file of chain 2 of ttr tetramer
   - posre_Protein_chain_C.itp : Position restraint file of chain 3 of ttr tetramer
   - posre_Protein_chain_D.itp : Position restraint file of chain 4 of ttr tetramer
   - topol_Protein_chain_E.itp : Topology file of the viral fragment (fi10, sk9, or hv11)
   - posre_Protein_chain_E.itp : Position restraint file of viral fragment (fi10, sk9, or hv11)

   Note that topol_Protein_chain_E.itp and posre_Protein_chain_E.itp are not available in control simulation directories since we performed our control simulations without any viral fragments.

3. md_parameter_files : Contains md parameter files used in our simulations.
    a. ions.mdp : Parameter file for ions addition
    b. minim.mdp : Parameter file for energy minimization
    c. nvt.mdp : Parameter file for NVT equilibration
    d. npt.mdp : Parameter file for NPT equilibration
    e. md.mdp : Parameter file for production MD run
